# Predicting characterization of microbiome taxonomy from imaging using machine learning approaches

**DOI:** 10.1101/2025.02.03.636311

**Authors:** Brandon Niese, Philip Sweet, Bryan Dinh, Amanda N. Scholes, Danielle S. LeSassier, Krista L. Ternus, Lydia Contreras, Vernita Gordon

**Affiliations:** Department of Physics, Center for Nonlinear Dynamics, The University of Texas at Austin, Austin, Texas, USA; Interdisciplinary Life Sciences Graduate Program, Department of Molecular Biosciences, The University of Texas at Austin, Austin, Texas, USA; Signature Science, LLC, 8501 North Mopac Expressway, Suite 100, Austin TX 78759; McKetta Department of Chemical Engineering, The University of Texas at Austin, Austin, Texas, USA; LaMontagne Center for Infectious Disease, The University of Texas at Austin, Austin, Texas, USA

## Abstract

For this study, a total of 47 mock human skin microbiome communities were created using microorganisms collected from human donors and grown *in vitro* for between eight and 32 days. Each mock community sample was split. Ten mL of each sample was used to determine the taxonomy of the community, using metatranscriptomics and Kraken2 to provide population-level taxonomic information; five mL of each sample was used for imaging. The resulting micrographs served as the basis for establishing a new analysis pipeline that sequentially used two different methods for machine learning and one statistical technique: (1) confocal microscopy images were segmented into individual cells using the generalist, deep learning, publicly available machine learning model Cellpose; (2) continuous probability density functions describing the joint distribution of the cell area and eccentricity were found using algorithms expressing the statistical technique of kernel density estimation; (3) these probability density functions were used as input for convolutional neural networks, that were trained to predict both the taxonomic diversity and the most common bacterial class, independently of metatranscriptomics. Specifically, models were made to predict the Shannon index (a quantitative measure of taxonomic diversity) and to predict the most common bacterial class, for each micrograph. Measured Shannon indices (based on metatranscriptomics) ranged from nearly 0 to 1.4. The model predictions of Shannon indices had a mean squared error of 0.0321 +/- 0.0035. The model predictions of the most common taxonomic class of bacteria had an accuracy of 94.0% +/- 0.7%.

**Importance:** Taxonomic diversity is a useful metric for describing microbial communities and can be used as a measure of ecosystems’ health, resilience, and biological interactions. Characterization of microbial community diversity also has diagnostic applications. For the human skin microbiome in particular, microbial diversity directly impacts skin health, including resilience against pathogens and regulation of immune responses. Currently, microbial diversity can be determined either using traditional staining methods that are limited to pure cultures or using sequencing methods that require high investment in cost, time, and expertise. In this study, we demonstrate an innovative method that employs microscopy images of bacterial communities and machine learning to predict taxonomic diversity and the dominant bacterial classes of bacterial communities. The underlying framework of the pipeline for taxonomy prediction has the potential to be adapted and extended to other organisms and microbiomes and to make taxonomic analyses less expensive and more feasible in low-resource settings.

## Introduction

Taxonomical metrics can be used to provide insights into the composition, diversity, and function of microbial communities; these metrics are key to understanding ecosystem dynamics, human health, and environmental processes [1–4]. For example, the Shannon index is a quantitative measure of taxonomic diversity that accounts for both the number of different types present and their relative abundance [5, 6]. Taxonomy can aid in identifying potential pathogens, understanding microbial interactions, and developing strategies for managing diseases [1, 2]. Additionally, knowledge of microbial taxonomy is essential for understanding how microbial communities contribute to ecosystem processes such as nutrient cycling and decomposition, and for identifying microbes with biotechnological potential [1–4].

An example of the importance of microbial taxonomy is the human skin microbiome. The human skin microbiome plays a crucial role in maintaining skin health and immune function [7]. Changes in the skin microbiome have been associated with disease [8–12]. Beyond the skin alone, changes in taxonomic composition and diversity of the microbiome are associated with changes from health to disease, and vice versa, in sites including the vagina, the eye, the ear, the nasopharynx, the mouth, and the gut [13, 14]. Microbial diversity also matters in many other contexts, including soil, plants, and waterways [15–22]. Thus, knowledge of the diversity of microbiomes has the potential to contribute to fundamental understanding of complex ecosystems and to lead to better health, for humans, animals, plants, and entire ecosystems. The ability to do rapid, facile assessments of microbiome taxonomy even in resource-limited settings would be a big step toward realizing this potential.

Multiple sequencing-based approaches exist to determine the taxonomy profile of a microbial community. Amplicon sequencing approaches, such as 16S sequencing, rely on targeting a specific genome region and then comparing differences in that sequencing to determine the species present. Metagenomic approaches, on the other hand, rely on sequencing the entire genome and assembling genomes from those reads or assigning taxonomy classifications to each read [23]. Finally, metatranscriptomics approaches rely on sequencing the transcriptome of the samples, generally after rRNA depletion, and then assigning a taxonomy classification to each read [24]. Comparisons between these methods have shown that metagenomics approaches capture a greater diversity of species than amplicon sequencing [25] or metatranscriptomics [26]. However, metatranscriptomics has been suggested to better capture which cells are metabolically active in a community, whereas metagenomics equally captures active and inactive cells [26].

However, several factors can make determining microbial taxonomy can be challenging, expensive, and time-consuming. Microbial communities can be highly diverse, with many different types of microbes present in a single sample, which makes their identification and classification complex. For example, the human gut can contain more than 1,000 different species of bacteria from at least ten different taxonomic families and the human oral microbiome can comprise members of at least six different taxonomic families [27]. Sample processing, including extraction of nucleic acids and preparation for sequencing, can be labor-intensive. Furthermore, the costs associated with next-generation sequencing technologies can be high (hundreds of United States dollars for each sample), particularly when analyzing large numbers of samples or when conducting metagenomic studies. Finally, the analysis of sequencing data and assignment of taxonomic labels require specialized bioinformatics tools and expertise, which can be time-consuming and computationally intensive.

For all these reasons, sequencing-based taxonomic analysis may not be feasible for all investigators, particularly for those in resource-limited settings. Moreover, advances in microscopy, data storage, and computational power have made imaging and machine learning (ML) accessible to researchers in resource-limited settings [28, 29]. For example, portable paper-based “foldoscopes” cost less than $100 with accessories, can magnify up to 340×, and allow images to be captured on a standard smart-phone microscope [29]. “Foldoscopes” are compatible with both Gram staining and dark-field microscopy, both of which can visualize bacteria. At a slightly higher price point of less than $1,000, portable open-source devices can image bacteria using fluorescence [30, 31]. Imaging technologies, particularly fluorescent microscopy, have enabled detailed visualization of microbial communities as well as measurements of biological activity such as metabolism and the production and distribution of proteins [32–34]. In this study, our goal is to develop and demonstrate a method for performing taxonomic analysis of multispecies communities using microscopy combined with ML. For this proof-of-concept demonstration, we used a straightforward amphiphilic fluorophore stain to visualize the bacterial cell envelope, which is the boundary defining cell morphology. Thus, all the microscopy-based taxonomic analysis presented here is based on measurements of cell morphology.

An imaging-based taxonomy approach that is widely applicable to microbiomes requires the ability to capture, with spatial resolution comparable to the size of the microbes involved, a wide range of micro-organism types in a single image [33, 34]. Furthermore, a pre-requisite for any microscopy-based taxonomic analysis of a microbial population is the ability to identify each individual cell in an image containing many cells. This process is known as “cell segmentation” and ML techniques are widely employed for this [35, 36]. Another form of ML, the convolutional neural network (CNN), has been trained to identify images of different types of bacteria via large-scale meta-analyses of image databases [37, 38]. CNNs excel in tasks such as computer vision and image classification because CNNs can extract features and reduce the dimensionality of images, decreasing their complexity, without losing important characteristics of the input [39]. A range of different types of artificial neural networks, including CNNs, have been used to extract, from micrographs, information about individual bacterial cells – information extracted includes identification of specific cell phenotypes and morphologies [40–42]. Thus, both cell segmentation and CNNs have well-established utility for analyzing images of bacteria and other cells.

However, images contain information on the spatial arrangements of cells that is not relevant for the type of taxonomic analysis we do here and consequently increases the computational burden to no benefit. To reduce the computational burden from that associated with direct CNN analysis of images, we use only morphological information obtained after cell segmentation. This morphological information is a set of discrete measurements, which we therefore need to re-cast into a continuous “image-like” form appropriate for CNN training. For this, we use the statistical technique of kernel density estimation (KDE) on discrete measurements to create a continuous probability density function (PDF) for the properties measured [43–45]. Since an image consists of one or more approximately continuous (modulo pixels) greyscale 2D maps, KDE is not typically appropriate for direct analysis of images. However, KDE can be used to obtain the probability distribution of the cellular characteristics measured using microscopy [46–48].

The resulting pipeline achieves taxonomical characterization of microbial populations from confocal microscopy images. Our approach is to sequentially combine techniques for: (i) ML segmentation to identify individual cells so that each cell’s area and eccentricity can be measured; (ii) KDE to find normalized PDFs that describe the bacterial population in terms of cell area and eccentricity; and (iii) ML by CNNs to identify the Shannon index and the most populous taxonomic class present.

A key feature of our work is that here we first process the images to produce PDFs and then apply neural networks for population analysis. Previous work has typically trained neural networks directly on images of one to a very few cells. The computational expense and time required for training CNNs on images scales at least linearly, and usually faster than linearly, with the number of pixels in the image; our images, each containing at least 100 bacteria, would be time-consuming and costly to use conventionally for CNN training. We reduce the demands of CNN training by discarding information on the spatial arrangements of cells so that only morphological information is used for training.

The workflow pipeline we present here has the potential to be adapted and extended to analyze other organisms and microbiomes, and to include subcellular structures, such as the nucleoid, as well as other, non-morphological measures of biological activity which could also be highly relevant to taxonomic analysis. It also has the potential to make taxonomic analysis of microbial communities faster and less costly than traditional approaches.

## Results

### Complex bacteria communities were derived from human donors

Complex communities of bacteria representative of the human skin microbiome were grown on Vitro-Skin® (VS) using microbial samples collected from human donors (**Figure 1**, details in methods). VS, an artificial material that is designed to mimic the characteristics of human skin, has been used in a range of *in vitro* investigations of scenarios in which skin plays a central role [49]; such scenarios have included pathogen transmission, protection against solar radiation, adhesion of medical devices, and spreading of emollient moisturizers [49–54]. After collecting microbes from the skin of ten human donors, the samples from all ten donors were pooled. This pooled suspension of microbes was then distributed into 50 initially identical cultures that were inoculated on VS. Samples were collected after 8, 11, 17 and 32 days of growth; bacterial communities were visible on the VS on all days. (Samples were also irradiated after seven days of growth; this was shown not to be relevant to these studies (**Supplementary Figure S1**)). At the time of collection after growth, each culture was divided, with five mL used for imaging and ten mL used for the extraction of nucleic acids for sequencing and taxonomic analysis. Out of 50 initial cultures, metatranscriptomics information was successfully obtained for 47. These 47 communities are the ones examined in the rest of this study.

**Figure 1.**
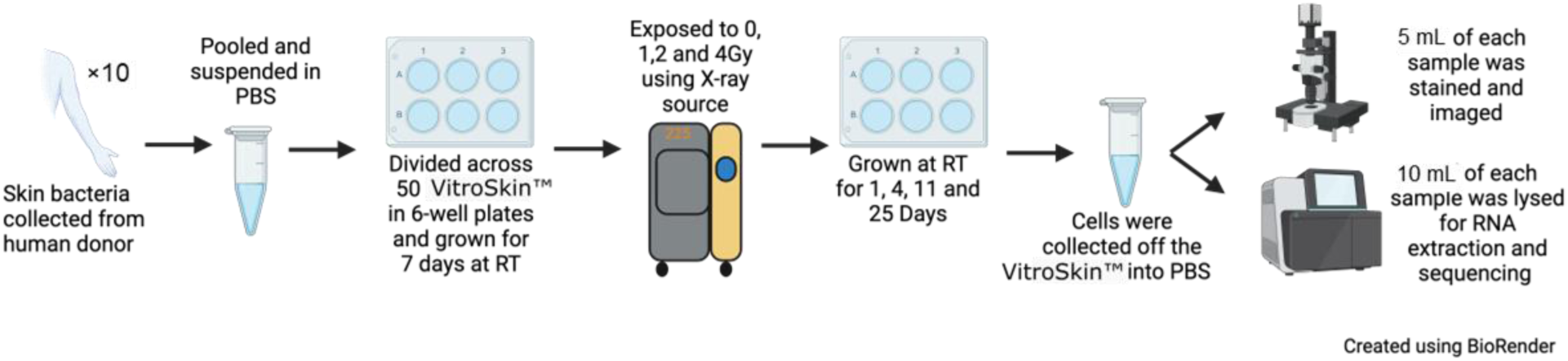
Methodology for generating skin microbiome mock communities, using VS, for RNA sequencing and imaging. Microbes were collected from the skin of ten human donors, pooled together, and this pooled suspension was used to inoculate 50 cultures on VS. Samples were irradiated after seven days of growth and collected after 8, 11, 18 and 32 days of growth following the initial inoculation; irradiation was shown to be irrelevant for these studies. At time of collection, all VS cultures were divided, with 5 mL used for imaging and 10 mL used for the extraction of nucleic acids for metatranscriptomics and taxonomic analysis. Out of 50 initial cultures, metatranscriptomics information was successfully obtained for 47. These 47 communities are the ones examined in the rest of the manuscript.

Metatranscriptomics and the Kraken2 taxonomic classification pipeline were used to determine the taxonomic makeup of the cultures grown on VS (**Figure 2**). Kraken2 assigns a taxonomic lineage to each RNAseq read by matching each k-mer in a query sequence to the lowest common ancestor of all reference genomes in its database that contain the k-mer. The final taxonomic label is assigned based on how similar the k-mer content of the sequence is to the k-mer content of the reference genome. While Kraken2 will provide classifications for reads at the species or strain levels, performance becomes more accurate at higher taxonomic levels [55–59]. Therefore, in this study we focused on the classes of microbes within the VS cultures. Differences in class-level diversity of microbiomes are associated with disease in cases including ulcerative colitis, mastitis, diabetes, and chronic obstructive pulmonary disease [56–59]. For each sample, the reads mapping to each class were determined. The top five most abundant classes (by fraction of reads from Kraken2 output) are presented in **Figure 2**. On Day 8 *Bacilli* was the most-abundant class in 90% of samples; on Day 11 either *Alphaproteobacteria* or *Bacilli* was the most-abundant class in 83% of samples; on Day 18 and Day 32 *Actinobacteria* was the most-abundant class in 65% of samples.

**Figure 2.**
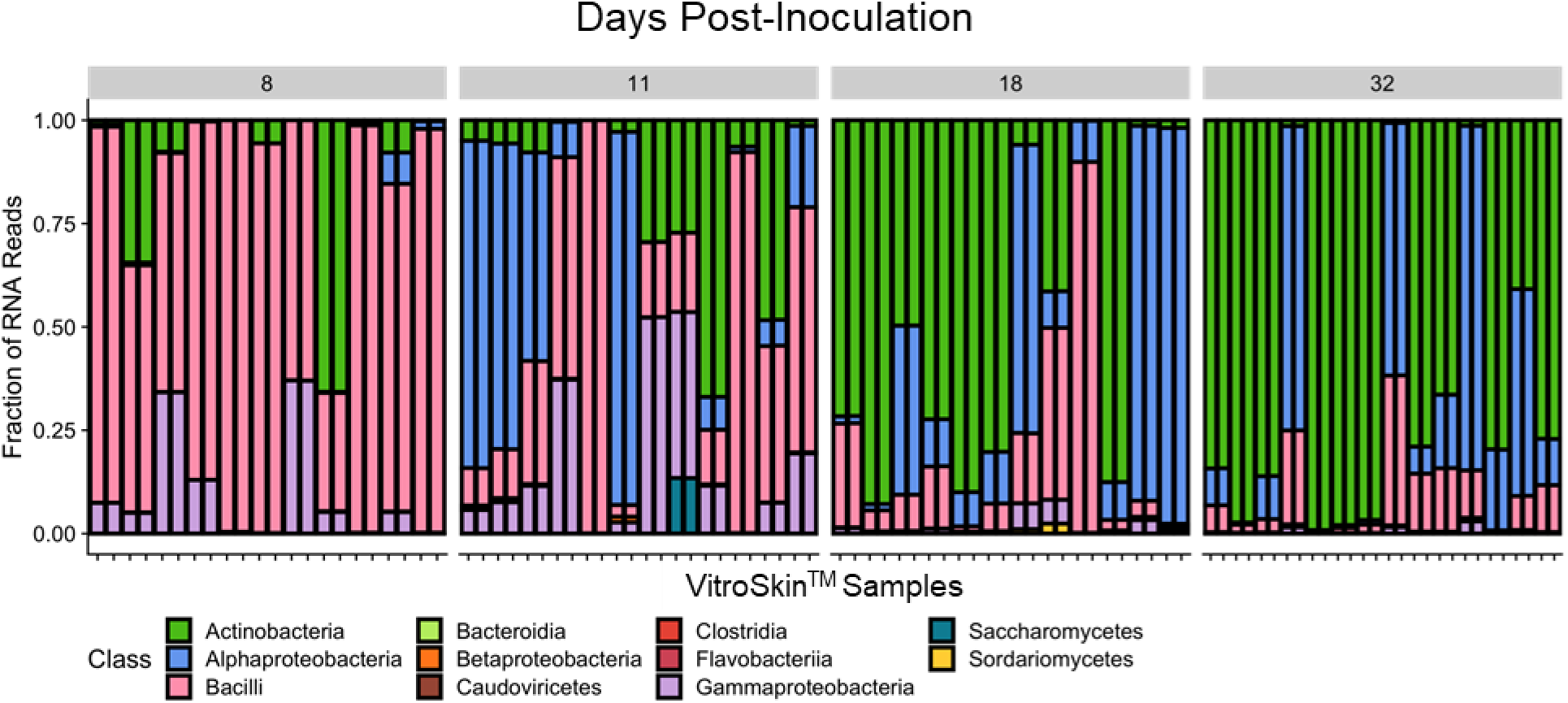
Taxonomy of cultures grown on VS. Class-level diversity of 47 cultures grown on VS, for each measurement timepoint, generated using Kraken2.

Overall, this analysis confirmed that the most common classes found for cultures grown on VS are *Bacilli*, *Actinobacteria*, *Alphaproteobacteria*, and *Gammaproteobacteria* (**Figure 2**). These are the same classes that other researchers have identified as being well-represented in bacterial samples collected from dry skin [7, 60, 61].

To further assess the degree to which the communities grown on VS are comparable to the actual human skin microbiome from which they were derived, we compared the taxonomy results from these samples (**Figure 2**) with taxonomy data that others have reported from patients with atopic dermatitis (AD) [62] (**Supplementary Figure S2**). Previous researchers collected samples of the skin microbiome directly from human patients and immediately extracted bacterial DNA for sequencing (e.g. without the *in vitro* culturing and irradiation steps present in this study for the mock communities) [62]. We use these previously collected data as a baseline for taxonomic characteristics that should be expected for the human skin microbiome. For each patient, the microbiome was sampled from both the site of an AD lesion and from adjacent, non-lesional, skin [62]. Thus, these data also provide a test case for the connection between microbial diversity and a healthy or diseased state at the site where the microbiome was sampled. The earlier researchers did a species-level taxonomic analysis [62]. We re-cast this information to class-level taxonomic analysis. Consistent with our findings for cultures grown in VS (**Figure 2**), the dominant classes within the samples from AD patients, from both sites with AD lesions (diseased) and without AD lesions (healthy), were *Actinobacteria* and *Bacilli*. Of the minor classes, *Gammaproteobacteria* were most common. Thus, the taxonomic classes found in bacterial samples cultured on VS were similar to those found in bacterial samples taken directly from human skin.

Besides the taxonomic groups present, another important property of microbial communities is the number and the relative abundance of species in a community. Alpha-diversity describes local diversity, within a single ecological site, and therefore is the appropriate measure for both the cultures grown on individual VS coupons and the samples taken from individual skin sites [63]. We quantify class-level alpha diversity using the Shannon index [5, 6]. The Shannon index *H*^′^ is given by

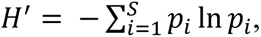

where *p*_*i*_ is the fraction of the sample population that is class *i*, and *S* is the total number of classes present in the sample population; ln is the natural logarithm. *H*′ has its lowest possible value of zero when the population is composed of a single class. As the diversity of class types in the population grows, *H*′ increases. Other researchers previously found that skin microbiome samples from AD patients had lower species-level Shannon indices for samples from lesional sites than for samples from non-lesional (healthy) sites [64]. We find the same trend for class-level Shannon indices (**Table 1**). We also found that the ranges of Shannon indices for mock communities grown on VS and for samples from AD patients had a percent difference of less than 10% (1.78 - 0.3 = 1.48 for AD patients and 1.44 - 0.07 = 1.37 for VS communities) (**Table 1**).

**Table 1:** Shannon Alpha-Diversity of Vitro-Skin®and Atopic Dermatitis (AD) samples. Shannon indices (mean, minimum, and maximum) for mock communities grown on VS (this study) and for samples taken directly from the skin of human patients with AD [63].

| Sample | Mean | Minimum | Maximum | Variance |
| --- | --- | --- | --- | --- |
| Vitro-Skin® | 0.69 | 0.07 | 1.44 | 0.12 |
| AD - Healthy | 1.07 | 0.09 | 1.70 | 0.25 |
| AD - Lesion | 0.84 | 0.03 | 1.78 | 0.09 |

A Welch’s t-test shows no statistically significant difference between the distribution of Shannon indices measured for VS communities and those found for lesional sites on AD patients (p = 0.08). In contrast, a Welch’s t-test comparing the distribution of Shannon indices of all the VS samples to the distribution of Shannon indices for non-lesional sites for AD patients gives a p-value of 5.298×10^-07^, indicating that these are different with statistical significance. Similarly, comparing the non-lesional sites to the non-monoculture VS samples yields a p-value of 0.003783, which also indicates statistically-significant difference.

Thus, the samples grown on VS show a reasonable approximation of the species and diversity that would be expected for a sample collected directly from human skin (specifically from lesional sites on AD patients). Moreover, the range of Shannon indices measured for VS samples is sufficiently large enough to encompass the diversity of directly-collected microbiome samples from both healthy and lesional sites for patients with AD.

Overall, in terms of both the classes represented and the diversity as quantified with Shannon indices, our mock communities grown on VS have strong taxonomic resemblance to direct samples of the human skin microbiome. An exception is the case of obligate anaerobes (e.g. *Clostridia, Bacteroidia*, and *Fusobacteria*), since culturing on VS was done in the presence of oxygen. This omission is notable, since anaerobic taxa are over 20% of determined taxa in the human skin microbiome [65]. However, the purpose of the mock communities generated by growth on VS was to demonstrate the utility of a new imaging-based approach to microbial taxonomy. The taxonomies and classes represented are sufficiently diverse for this purpose.

Class-level taxonomic results for VS cultures and samples from AD patients can be found in the Texas Data Repository [66].

### Fluorescent microscopy of bacterial mock communities captures geometric properties of individual cells

For each of the 47 samples successfully sequenced (**Figure 1**), the portion of the microbial content that was not used for metatranscriptomics was imaged using spinning-disc confocal fluorescence microscopy. For this analysis, the cell membranes were stained using the fluorescent stain Nile Red, allowing us to visualize the cell envelope of each bacterium. Each confocal micrograph consists of a greyscale image of approximately 1,000 cells. A sample image is shown in **Figure 3A**. Each of the 47 samples was imaged in nine different fields of view. Of the resulting 423 images, three contained fewer than 100 cells and were not used for further analysis; these images all came from the same culture containing 99% *Actinobacteria*. The remaining 420 images were used for all image-based analysis; images from the same mock community were used as technical replicates for that community. All images used are available through the Texas Data Repository [67].

**Figure 3.**
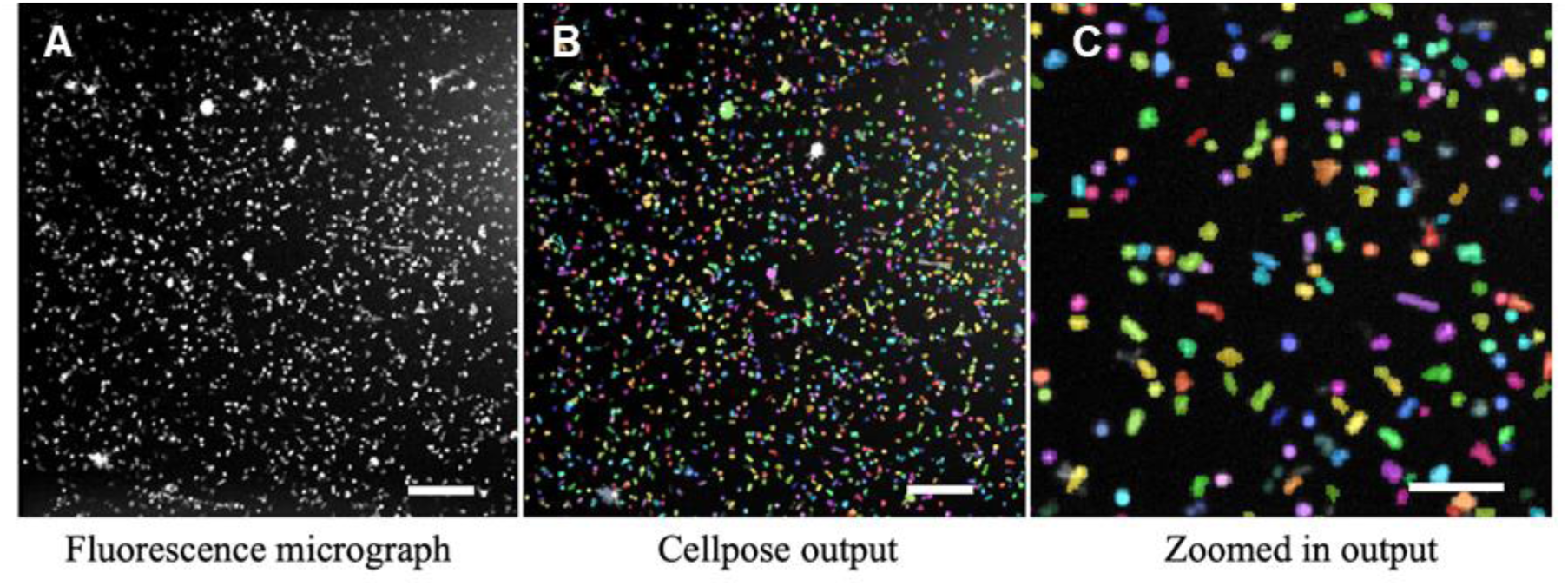
Imaging bacterial populations and segmenting individual cells. (A) Spinning-disc confocal fluorescence micrograph of a mock community grown on VS. Scale bar is 20 μm. (B) For the same image, output of the Cellpose segmentation algorithm. False colors indicate individual segmented cells. Scale bar is 20 μm. (C) Zoomed in section of the Cellpose output in panel B. Scale bar is 10 μm.

The images were then segmented into individual cells, using the ML segmentation library Cellpose [36]. Examples of segmented output are shown in **Figure 3B-C**, in which false coloring is used to distinguish different segmented cells. Segmented regions were used to create masks, with each mask showing the fluorescence “footprint” of one cell membrane. These masks were analyzed to measure geometric properties of cells. This approach is based on a method we recently developed for high-throughput quantitative analysis of micrographs of large populations of bacteria [68].

The size of each cell is measured as its cross-sectional cell area, found from the area covered by the mask. The shape of each cell is measured as its eccentricity, found by fitting an ellipse to each mask and calculating its eccentricity. Eccentricity can be used to distinguish rod-shaped bacteria from round cocci, and it can be used to distinguish longer rods from shorter rods. The eccentricity for a round bacterium will be close to zero, and the eccentricity for a highly elongated rod will be close to one.

The outcome of these measurements is, for each image, a list of bacterial cells, each with measurements for area and eccentricity.

### Joint probability distribution functions describe the properties of a microbial population in geometric parameter-space

We expect that cultures that are taxonomically similar will also have similar joint probability distributions of cell area A and cell eccentricity e. In terms of these morphological characteristics, the measurements described in the preceding section allow each bacterium to be represented as a discrete point in a two-dimensional parameter space defined by two variables, cell area and cell eccentricity. KDE is used to convert this discrete distribution into a continuous, normalized PDF of two continuous random variables (cell area *A* and cell eccentricity *e*). For *x* = [*A*, *e*], the joint PDF ^*f*^^(*x*) allows the probability density of geometric properties to be visualized as an “image” in a parameter space spanned by cell area and cell eccentricity.

Using the taxonomy data from RNA sequencing, we assigned images (and the corresponding PDFs) as coming from homogeneous populations (if the top taxonomic class contained over 90% of the total bacterial population) or heterogeneous populations (if the top taxonomic class contained less than 60% of the total bacterial population). Populations in which the top taxonomic class made up between 60% and 90% of the total bacterial population were considered neither homogeneous nor heterogeneous. Using these data, we assigned 96 images to homogeneous cultures and 99 images to heterogeneous cultures; 225 images were from cultures considered neither homogeneous nor heterogeneous. Of the 96 images assigned to homogenous cultures, 51 were mostly composed of *Actinobacteria*, nine were mostly composed of *Alphaproteobacteria,* and 36 were mostly composed of *Bacilli*. Since a given culture can be represented by up to nine images, this corresponds to seven homogeneous *Actinobacteria* cultures, two homogeneous *Alphaproteobacteria* cultures, five homogeneous *Bacilli* cultures, and ten heterogeneous cultures. Averaging together all PDFs for each of these types of culture shows consistent differences between the four population categories **(Figure 4**).

**Figure 4.**
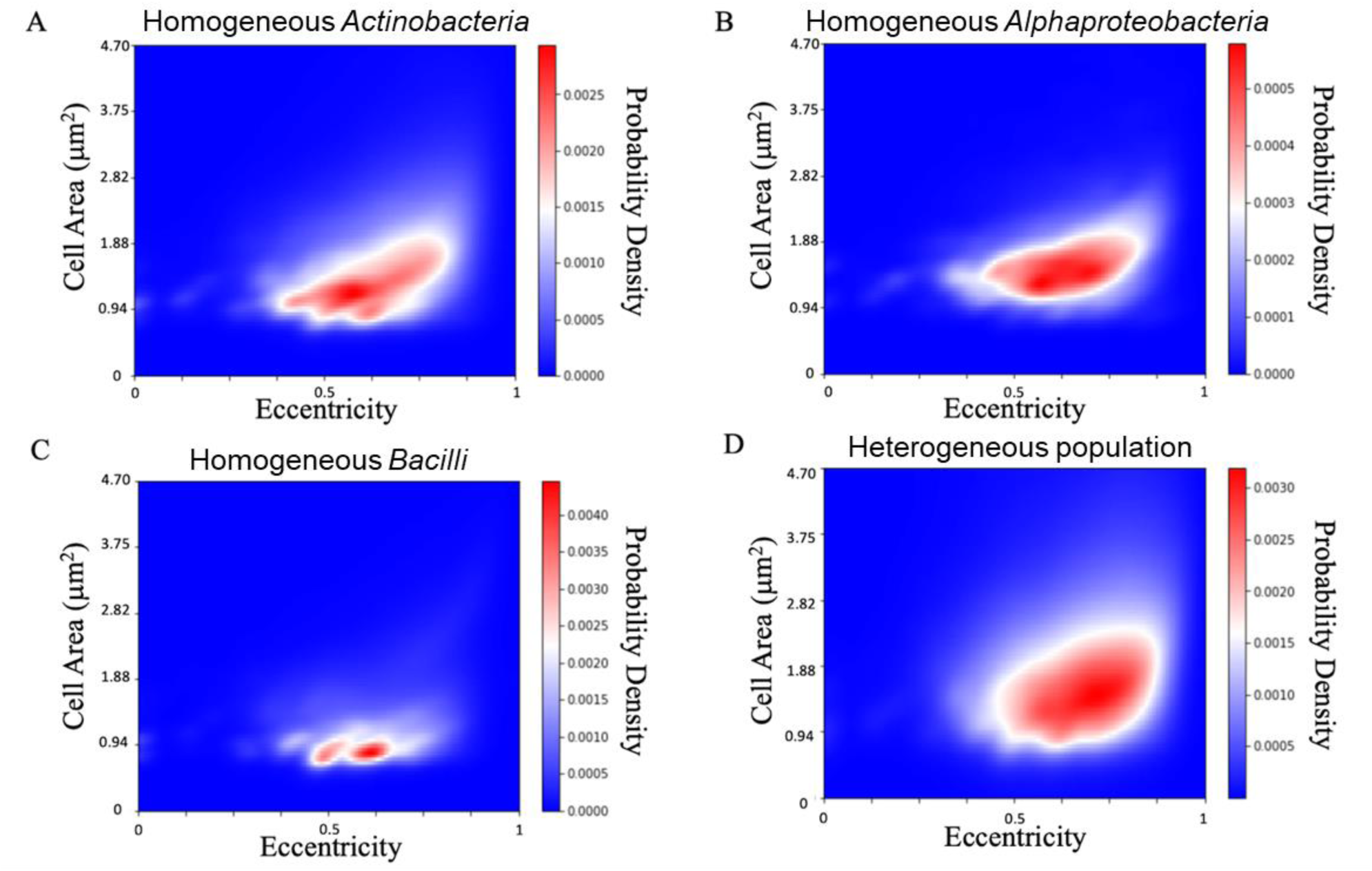
2D Probability Density Functions describing four types of bacterial populations. The normalized PDF can be thought of as a 2D histogram that describes the distribution of morphologies in a population. (A) Average PDF plot of bacterial populations with a top percentage of *Actinobacteria* greater than 90 percent (average of 7 cultures). (B) Average PDF plot of bacterial populations with a top percentage of *Alphaproteobacteria* greater than 90 percent (average of 2 cultures). (C) Average PDF plot of bacterial populations with a top percentage of *Bacilli* greater than 90 percent (average of 5 cultures). (D) Average PDF plot of bacterial populations with any top-class percentage of less than 60 percent (average of 10 cultures).

To further assess the PDFs for possible connections with taxonomy, we compare each culture’s average PDF with every other culture’s average PDF. This provided comparisons across cultures with varying levels of taxonomic diversity and different top taxonomic classes. In every case, the PDFs obtained from each individual image from a culture were averaged to get an average PDF describing that culture. Two example PDFs of individual cultures are shown in **Supplementary Figure S3A and S3B**. Plots of all PDFs generated are available through the Texas Data Repository [67].

To compare two cultures’ PDFs and test our hypothesis that cultures that are taxonomically similar will also have similar joint probability distributions of cell area and cell eccentricity, we subtract PDFs and take the absolute value of the result. An example of the absolute values of the differences between pairs of PDFs is shown in **Supplemental Figure S3C**. To consolidate the morphological difference between two cultures into a single value, we calculate the numerical integral of the absolute value of the function found by subtracting, element-wise, the two cultures’ PDFs. This process was repeated for every pair of cultures grown on VS (**Supplementary Figure S4**). We term the absolute value of the difference between two cultures’ PDFs the morphological difference, and we expect this value to be small (near zero) for very similar populations and large (near two) for very different populations.

Importantly, from this analysis we find that the average morphological difference between samples with the same top taxonomic class is significantly lower than the average difference between samples with different top taxonomic classes (**Supplementary Figure S5**). This supports the notion that populations that are taxonomically similar also have similar population distributions in the morphological parameter space defined by eccentricity and cell area.

### Predicting Shannon index using a convolutional neural network (CNN)

We next trained a CNN to predict the Shannon index of a given PDF so that we can assess the class-level diversity of a sample using morophological characteristics obtained from microscopy. For this, we used the Adam optimization algorithm, which optimizes using first-order gradient descent [69]. This algorithm leverages the large pre-existing body of work on ML for image recognition [70–75], is computationally efficient, and is suited for large parameter spaces [69]. The loss function optimized by the Adam algorithm is the mean squared error (MSE) of predicted versus true values for the Shannon index. To train this model, the total set of all 420 PDFs obtained from 420 images is split into a training set and a testing set. The training set is composed of 70 percent of the 420 total PDFs, i.e. PDFs representing 294 images from randomly selected cultures and fields of view. The testing set is composed of 30 percent of the 420 total PDFs, i.e. PDFs representing 126 images from randomly selected cultures and fields of view.

For k = 10 iterations, the data are all randomly split anew into training and testing sets [76, 77]. One complete pass of the training set through the CNN algorithm is termed an epoch. After 70 epochs of training, the average performance metric for each train/test set is calculated to assess the model’s accuracy and averaged over the k = 10 train/test splits. We used MSE as the performance metric. After 70 epochs, the average MSE of the training sets was 0.00477 (+/- 0.00072, standard error of the mean) and the average MSE of the testing sets was 0.0321 (+/- 0.0035, standard error of the mean) (**Supplemental Figure S6A**). Thus, the MSE of the testing sets is an order of magnitude smaller than the mean value for the Shannon index, 0.69, and two orders of magnitude smaller than the size of the full range of Shannon indices measured, 1.37 (**Table 1**). Most notably, the MSE of the testing sets is one order of magnitude smaller than the variance in Shannon indices measured for all VS samples (**Table 1**). If the model’s prediction for Shannon index was always the mean value, then the MSE would be exactly equal to the variance. Always predicting the mean value corresponds to random guessing; thus, this model’s MSE is an order of magnitude smaller than would be obtained by random guessing and the model’s accuracy is an order of magnitude better than would be obtained by random guessing.

**Figure 5** shows the plot of predicted versus true Shannon index of each PDF for one of the training-testing data splits used. From this, we conclude that the model developed by machine learning is a reasonable predictor of the taxonomic diversity of these samples from data obtained from micrographs of the samples.

**Figure 5.**
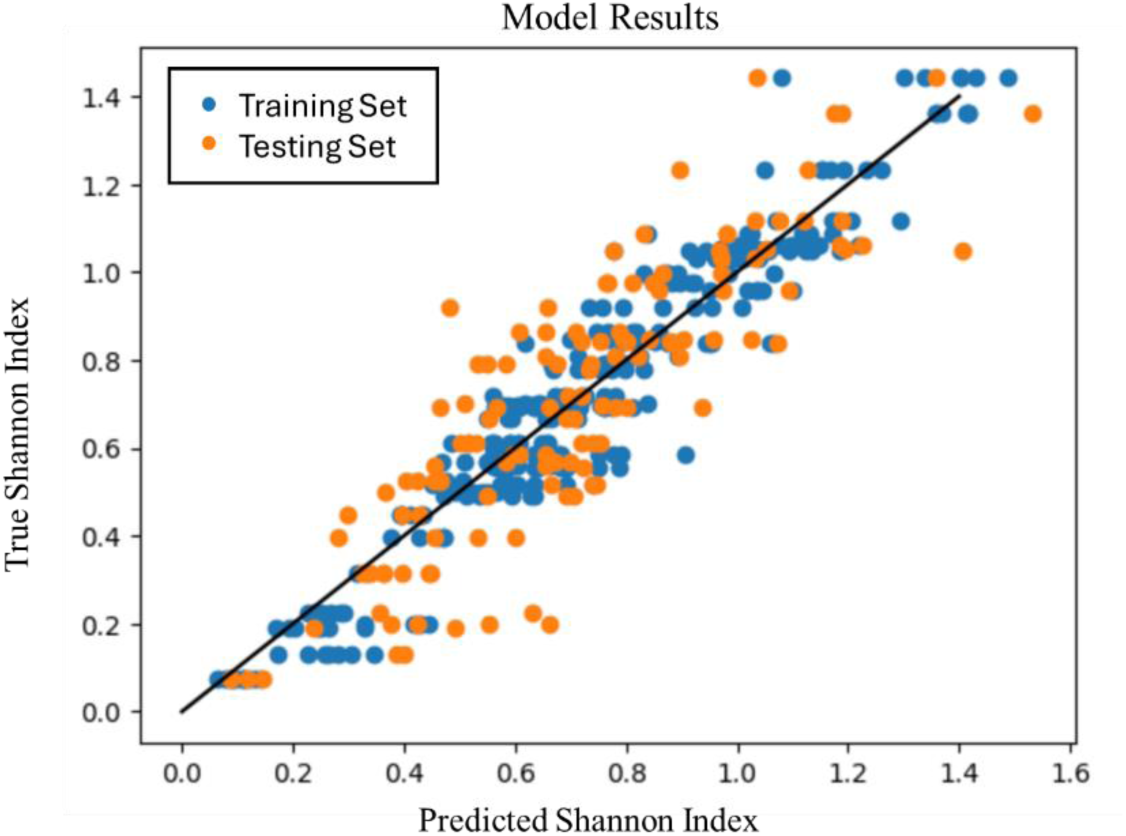
Results of the model for predicting Shannon index. Each dot indicates for one sample (on the vertical axis) the true (from RNA sequencing) Shannon index and (on the horizontal axis) the predicted (from the CNN model) Shannon index. Blue dots indicate values for each sample in a training set of 294 samples. Orange dots indicate values for each sample in the corresponding testing set of 126 samples.

### Predicting the top taxonomic class using a CNN

We also developed a second CNN to predict the most abundant taxonomic class from a PDF representing a particular image. The CNN model for predicting the most abundant taxonomic class was trained using a 70/30 train/test split and repeated k-fold cross validation with k = 10, as for the CNN model for predicting the Shannon index, above. We find that the accuracy of the training sets is 100% +/- 0% and the accuracy of the testing set is 94.0% +/- 0.7% after 70 epochs (**Supplemental Figure S6B**).

This model was optimized using the same Adam algorithm as the model for predicting the Shannon index, above [69]. However, the loss function for training this model was sparse categorical cross entropy, which measures the difference between two probability distributions [78]. In the context of ML, cross entropy is often used as a loss function to quantify the difference between the predicted probability distribution and the actual distribution of the labels in a classification problem [79, 80]. It calculates the difference between the predicted probabilities and the true distribution [78]. The formula for sparse categorical cross-entropy is

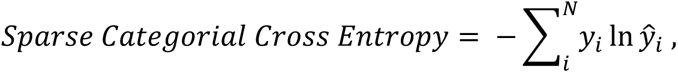

where y_i_ is the true probability distribution and ŷ_i_ is the predicted probability distribution for class [78]. The true labels are represented as integers indicating the class [81, 82].

**Figure 6** shows the confusion matrix of a testing set after 70 epochs of training. Therefore, we conclude that this CNN can accurately identify the taxonomic class of bacteria using information that originated with micrographs of the samples.

**Figure 6.**
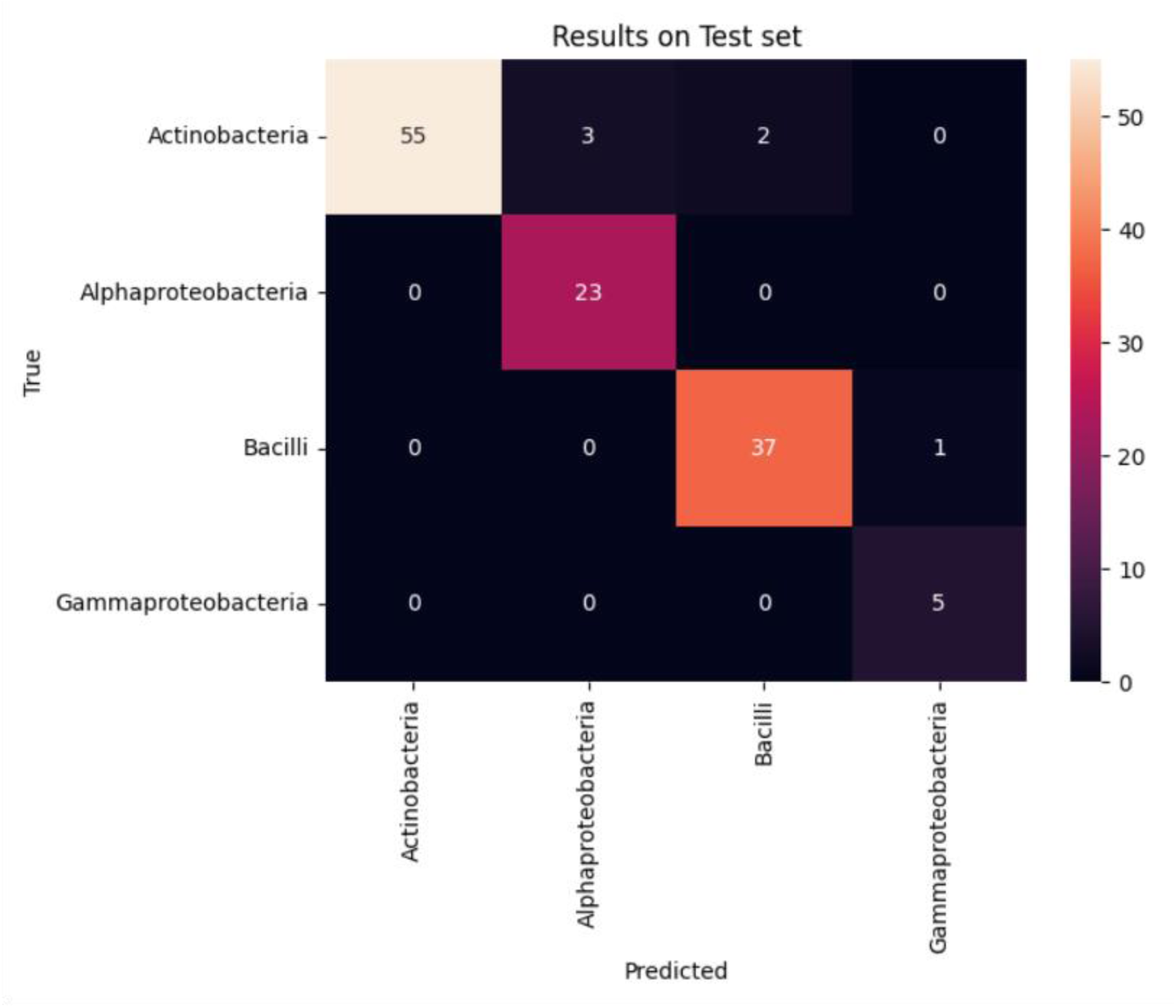
Accuracy of the model to predict the dominant taxonomic class, after 70 epochs of training, shown as the confusion matrix of the testing set. The accuracy of the corresponding training set was 100% and the accuracy of testing set was 94%. The columns represent the predicted values of the model, and the rows correspond to the true values of the corresponding data. 55 PDFs were accurately predicted to correspond to *Actinobacteria*-dominated cultures. 26 PDFs were predicted to correspond to *Alphaproteobacteria*-dominated cultures. Of those 26 samples, 23 were in fact dominantly *Alphaproteobacteria* as determined from sequencing, while three were mislabeled *Actinobacteria*-dominant samples. 39 PDFs were predicted to correspond to *Bacilli*-dominated cultures. Of these, 37 were indeed *Bacilli*-dominated, and two were mislabeled *Actinobacteria*-dominated samples. Six PDFs were predicted to correspond to *Gammaproteobacteria*-dominated cultures; five of these predictions were accurate, and one was a mislabeled *Bacilli*-dominated culture.

## Discussion

In this work, we used mock bacterial communities that we generated from initial collection from human skin. We validated these mock communities as reasonable models for the human skin microbiome by comparing with the published work of others [63]. Using metatranscriptomics, we find that our mock communities are comparable with microbial samples taken directly from human skin in both their class-level taxonomic diversity, measured using the Shannon index, and their dominant taxonomic classes (except for obligate anaerobes) (**Fig. 2**). We expected that after multiple days of culturing on VitroSkinTM that the bacterial community would have a significant number of dead cells. By washing the cells before imaging and using a dye for intact membranes, we selected live cells before imaging. To align the taxonomy data with the cells captured during imaging and then use it for model training, we utilized metatranscriptomics, which is biased toward metabolically active cells [26]. Using our mock bacterial communities, we demonstrate an approach to using ML for taxonomic analysis of bacterial populations containing multiple classes. The starting point for this approach is imaging of bacterial populations. Then, we use a sequential pipeline of computational tools. First, we segment images to identify individual cells so that their morphological properties can be measured. Second, we estimate the joint PDFs describing the likelihood that a cell in an image will have a given combination of values for its morphological properties. Finally, we train CNNs to predict the dominant taxonomic class and class-level diversity from the PDFs describing the morphological properties of cells in a population (**Fig. 7**). This multi-tool pipeline offers a significant reduction in both time and monetary costs compared to RNA sequencing (hundreds of dollars to sequence a sample versus a couple of minutes of training time on a mid-range laptop). The ability to use ML to gain insight into the taxonomy of populations, with modest computational investment (i.e. a 2023 MacBook Pro, with 16 GB of memory, and an Apple M2 Pro processor), paves the way for wider surveys covering more samples than would be tractable using sequencing.

**Figure 7.**
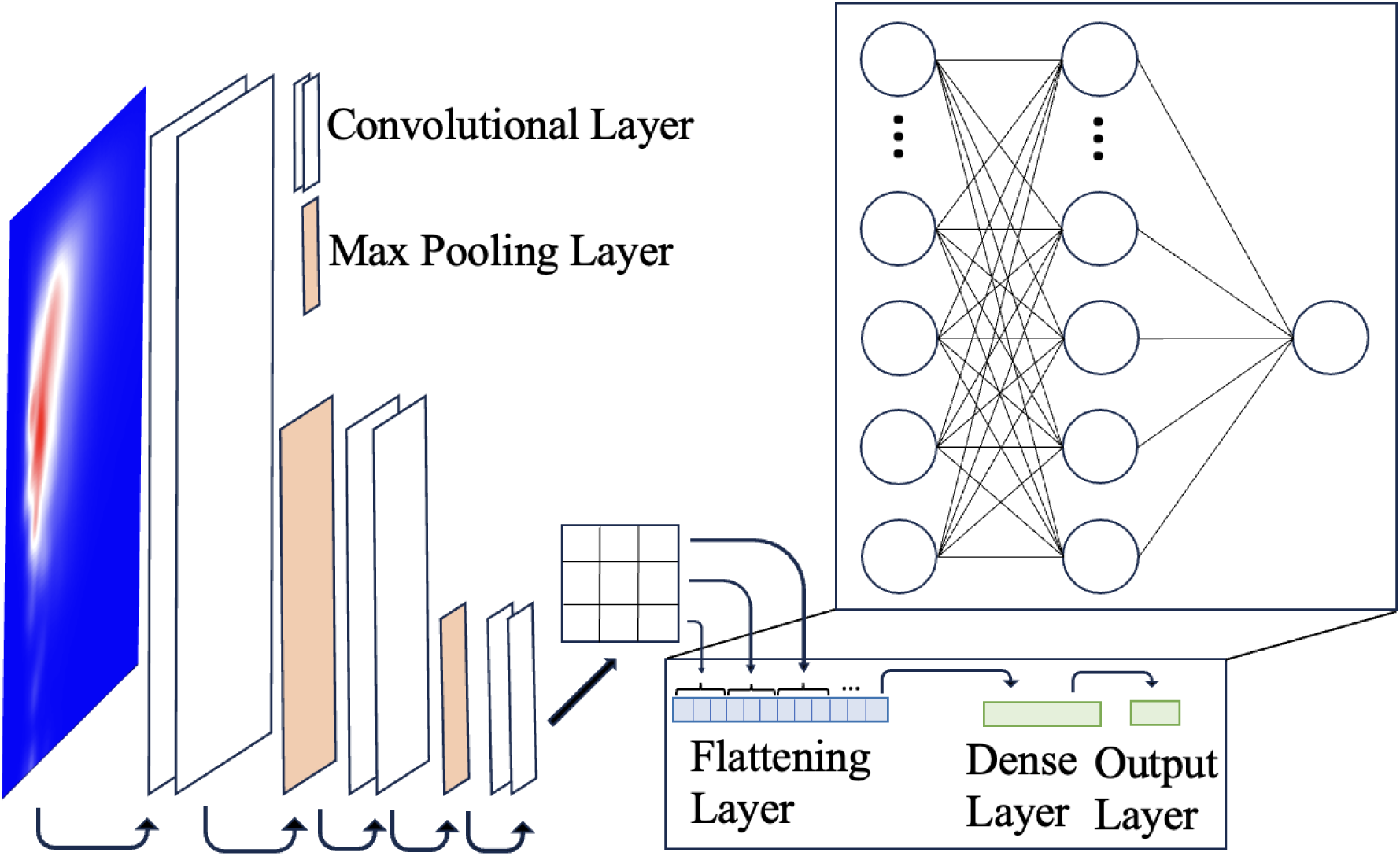
Schematic of the Convolutional Neural Network (CNN) architecture used in this study. The network comprises an input layer followed by a sequence of convolutional layers interspersed with max-pooling layers. Subsequently, a flattening layer is employed to transform the output of the convolutional layers into a one-dimensional array, which is then fed into a dense layer. The final output layer produces the predicted results.

The choice of pre-processing used to go from micrographs to the inputs for the CNN ML models was important for facilitating rapid training of the CNN models on a mid-range laptop. Micrographs of bacterial population contain information not only on the morphology of individual cells, but also the position and orientation of each cell with respect to every other cell in the image. Although the richness of spatial information provided can be a strength of microscopy, the position and orientation of bacteria are not relevant for the taxonomic analyses we do here. Therefore, to reduce the time required to train CNN models for taxonomic prediction from images, we reduce the amount of information fed to the CNNs by extracting only morphological information from micrographs. Here, each of the individual micrographs is 922 × 922 pixels, which means each micrograph corresponds to an array with 850,084 elements. In contrast, each PDF array is 200 × 200, or 40,000 elements. This value is more than an order of magnitude smaller than the number of elements in the micrograph from which the morphological information originated. The computational expense and time required for training CNNs on images scales at least linearly, and usually faster than linearly, with the number of elements. Therefore, this approach—extracting and using only morphological information, rather than the entire image—is an effective approach to speeding up CNN training by at least an order of magnitude and possibly much more.

Morphological properties other than cell size and cell eccentricity also have the potential to be extracted from micrographs. In our recent work, we also used microscopy to measure the size, circularity, and eccentricity of the nucleoid, and the relative size of the nucleoid with respect to the cell [68]. These measurements can all be extracted from micrographs of bacteria that have been segmented using Cellpose [68]; combining them together with cell size and cell eccentricity would allow determination of joint PDFs of a six-dimensional variable *x* = [*A*, *e*, *NA*, *Nc*, *Ne*, *NFA*]; this PDF ^*f*^^(*x*) would describes the likelihood of bacteria in a population having a cell area of *A*, a cell eccentricity of *e*, a nucleoid area of *NA*, a nucleoid circularity of *Nc*, a nucleoid eccentricity of *Ne*, and a nucleoid fractional area of *NFA*. Such an approach would allow a CNN to be trained to use more detailed morphological information while still avoiding the training time and cost associated with working with the original images and the extraneous positional and orientational information they contain. Inclusion of more morphological information has the potential to extend the number of classes that can be distinguished using this microscopy-based approach. Fluorescence microscopy can also provide measurements of biological activity such as metabolism and the production and distribution of proteins [32–34]. Measurements of this type of biological activity, obtained from the same imaging used to measure morphological characteristics, could also be incorporated into PDFs, to allow CNNs to be trained on a combination of morphological, proteomic, and metabolic data. The increased experimental capacity provided by this method could allow more taxonomic studies in situations where conventional sequencing-based methods are limited by the infrastructure or resources available.

To contextualize the results provided by our approach with a medically-relevant example, we recall our analysis of samples collected from lesional and non-lesional sites of patients with AD in a previous study [63]. The average class-level Shannon indices for AD samples from lesional and healthy sites are different by about 0.23 (**Table 1**). (While the AD study used amplicon sequencing to characterize taxonomy and in this paper we employed metatranscriptomics, comparative studies have shown that for abundant organisms, both amplification and metatranscriptomics approaches produce similar results [25, 26].) This difference is two orders of magnitude greater than the average MSE found for our testing sets in the CNN model to predict Shannon index. The MSE can be considered a measure of the CNN model’s resolution in predicting Shannon index. Both the lesional and the healthy sites are represented by a wide range of overlapping Shannon indices (Table 1); thus, given one sample from each type of site, a model such as ours would not be able to say which sample came from a lesional site and which from a healthy site. However, because the model’s resolution is much smaller than the difference in average Shannon index for each type of site, we expect that a model such as ours should be able to confidently detect overall differences in microbiome diversity between healthy and diseased sites, given a sufficient number of samples from each type of site. From this, we infer that an approach like the one we demonstrate here has potential to indicate whether there are differences in the microbiome associated with disease and the corresponding microbiome associated with health, if the number of patients or sites sampled for each were high enough.

### Conclusion

In conclusion, we have used taxonomic data obtained through RNA sequencing to train ML models to assess taxonomic diversity and top taxonomic class from imaging data alone. The model to assess diversity predicted the value of the Shannon index with a MSE that was within 5% of the average value of all the Shannon indices obtained through sequencing. The model to assess top taxonomic class did so with 95% accuracy. These findings were achieved on a mid-range laptop computer by using ML to do pre-processing of image data to be used to train a CNN. This scheme for rapid, lower-cost assessment of taxonomic properties has the potential to be applied to many different types of microbiomes and could be used to facilitate more broad-sweeping studies and studies.

## Acknowledgements

RNA sequencing and imaging by scanning-disc microscopy were done through core facilities at the University of Texas at Austin, namely the Microscopy and Flow Cytometry Facility (RRID:SCR_021756) and the Genome Sequencing and Analysis Facility (RRID:SCR_021713), which are overseen by the Center for Biomedical Research Support at the University of Texas at Austin. This work was supported by the Air Force Office of Scientific Research (Grant FA9550-20-1-0131) and the Welch Foundation (Grant F-1756). This research is also based upon work supported in part by the Office of the Director of National Intelligence (ODNI), Intelligence Advanced Research Projects Activity (IARPA), via Contract No. W911NF-22-C-0051. The views and conclusions contained herein are those of the authors and should not be interpreted as necessarily representing the official policies or endorsements, either expressed or implied, of the ODNI, IARPA, ARO, or the U.S. Government. We thank Prof. Scott Kravitz (Department of Physics, University of Texas at Austin) for very helpful discussions about machine learning. We also thank Madeleine Pont and Nicolette Keplinger (Signature Science, LLC) for support with human donor collections, Dr. Josh Gil (Signature Science, LLC) for assistance with formatting taxonomic summary files, and Drs. Myles Gardner and Curt Hewitt (Signature Science, LLC) for helpful technical discussions and manuscript review.

## Author Contributions

BN designed and implemented the data analysis and ML workflow for microscopy images and contributed to writing the paper. PS conceived of the project, contributed to taxonomic analysis of VS samples and published data from AD patients, and contributed to writing the paper. BD prepared samples for microscopy and collected the microscopy data. AS and DL developed the methodology and model for culturing on Vitro-Skin®, collected samples from human donors, and prepared microbiome samples. KT did RNA sequence analysis and bioinformatic taxonomic identification. LC and VG contributed to project design, to interpretation of data, and to writing the paper.

## Data availability

The authors confirm that the data supporting the findings of this study are available within the article and its supplementary materials. The code used to generate data sets from microscopy images and the code used to calculate taxonomic values from sequencing data can be found at https://github.com/bniese/TaxonomyProject. Taxonomic data, micrographs, and plots of PDFs for individual cultures can be found in the Texas Data Repository [66, 67].

## Materials and Methods

### Collecting and Culturing Samples

All human subject material was collected using protocols and informed consents that were approved by the Signature Science Institutional Review Board at Advarra under IRB protocol 00039337. Consent was obtained orally and in writing. Bacteria were collected from the forearms of human donors using flocked nylon swabs (Copan). Donors were excluded from the study if they had washed the collection area less than 10 hours prior to collection or if they had used oral or topical antibiotics in the last 72 hours, resulting in a total of 10 donors. Eight swabs were collected from each donor, four per forearm, with each swab collected from a new area of skin. Swabs were wetted with phosphate-buffered saline (PBS) with 0.1% Tween20 and swabbed across approximately a 3 × 3 cm area. Swab heads were snapped into Lyse&Spin baskets (Qiagen), covered with sterile PBS, and vortexed at 900 rpm at room temperature for 10 min. The sample was centrifuged through and transferred into a single collection tube. This was repeated for all donor swabs resulting in a pooled microbiome mixture. The pooled mixture was pelleted for 10 min at 5,000 × g and the liquid removed, careful not to disturb the pellet. The pellet was then resuspended in fresh, sterile PBS and kept on ice until inoculation. This approach was chosen to promote a greater diversity of bacteria types initially present in each culture. This suspension was divided across 50 hydrated Vitro-Skin® (IMS division of Florida Suncare Testing, Bunnell, Florida) coupons, so that each coupon was inoculated with 100 μL of the pooled microbial mixture. Inoculated Vitro-Skin® samples were dried and transferred to Reasoner’s 2A (R2A) agar plates, inoculated side facing up. Samples were supported on R2A agar and grown at room temperature in the dark for seven days. Vitro-Skin® coupons were then transferred to fresh R2A agar plates and irradiated using a Faxitron MultiRad 225 X-ray source (Wheeling, Illinois). Seven days after inoculation, radiation exposures were conducted at 200 kV and 12 mA using a 0.5 mm Al filter to select for high energy particles to 0, 1, 2 and 4 Gy. Twelve biological replicates were exposed at each dose, allowing for three biological replicates of each dose at each planned collection timepoint (1, 4, 11, and 25 days post-radiation corresponding to 8, 11, 18, and 32 days post-inoculation). Cells were kept at room temperature during the exposure. Cells were returned to growth chambers and incubated in the dark at room temperature, supported on R2A agar. On day 14 post-radiation, remaining samples were transferred to fresh R2A agar plates and continued incubating.

Cells were collected for morphology analysis at 1, 4, 11, and 25 days after exposure to radiation. To collect the microbial content off the Vitro-Skin®, the coupons were placed in 15 ml of PBS and vortexed for 3 min. The suspension was split into a 10 mL sample for RNA sequencing and 5 mL sample for imaging. The RNA-seq sample was used for sequencing to obtain taxonomic information, namely the proportions of each bacterial class present in every sample. The imaging sample was used for microscopy imaging. Both samples were pelleted and resuspended in either DNA/RNA Shield (Zymo) for sequencing or 10% formalin for imaging. The RNA-seq samples were extracted and cleaned up using the Quick-DNA/RNA MiniPrep Plus kit (Zymo) and Monarch RNA cleanup kit (NEB).

### RNA Sequencing

RNA was submitted to the Genome Sequencing and Analysis Facility at the University of Texas at Austin (https://wikis.utexas.edu/display/GSAF/Home+Page) for RNA quality analysis via Bioanalyzer 2100 (Agilent) and library preparation. Ribosomal RNA depletion was conducted using the RiboZero Plus kit (Illumina). The NEBNext Ultra II RNA kit (NEB) was used for library preparation. 150 base pair, paired-end (PE) sequencing was conducted across four lanes of a NovaSeq S4 chip (Illumina).

### Taxonomy Analysis

The FASTQ files obtained from Illumina RNA sequencing of the VS community samples were processed through the MetScale pipeline (https://github.com/signaturescience/metscale), which included generating visualizations and summaries of read quality with FastQC [83] and MultiQC [84], sequence read filtering and adapter trimming with Trimmomatic [85], and taxonomic classification on filtered Illumina paired-end reads with Kraken2 [55]. All complete genome or chromosome level assemblies from the NCBI RefSeq v.99 database for archaea, bacteria, protozoa, fungi, and viruses were downloaded and used for building a classification database for Kraken2 (k=35, ℓ=31). Kraken2 outputs were summarized with the bit-kraken2-to-taxon-summaries function from the bit package [86]. The MultiQC summary html report and the summarized Kraken2 classification results are provided through the Texas Data Repository [66]. For each sample, the percentage of reads mapping to each taxonomic class was determined from the Kraken2 outputs for further downstream processing. The Vegan R package was used to calculate the Shannon index [87]. Principal component analysis (PCA) was conducted on the number of reads assigned to each class within each sample using the R function pcrcomp() from the stats package [88]. Visualization was performed using the ggplot2 package [89]. PCA revealed no significant correlation between the taxonomic composition and the administered radiation doses, as indicated in **Supplemental Figure S1**.

### Microscopy

Confocal microscopy was performed using a Nikon W1 spinning disk confocal microscope (Nikon, Tokyo) using a Yokogawa W1 scanning head (Yokogawa, Sugar Land, Texas) with multiple pinholes for ultrafast confocal imaging. Imaging used a Plan Apochromat VC 60X water immersion objective in a 561 nm laser line.

### AD Samples

Previously published taxonomy data was acquired from [62]. The 16S counts matched to each taxonomy unit were tabulated in R, and the percent of reads mapping to each class.

### Staining Protocol

Cells were stained using 3 mM stock solution of Nile Red (Thermo Fisher, Invitrogen) (used for staining membranes). Stock solutions of Nile Red were made by dissolving solid Nile Red stain into DMSO at a concentration of 3 mM. A 100 μL of cell solution was aliquoted and 2 μL of stock Nile red solution was added. Cells were then placed in a rotary shaker for 30 minutes. Cells were centrifuged and the stain solution was removed. The cell pellet was then resuspended in 100 μL of Dulbecco’s Phosphate-Buffered Saline (DPBS). Cell suspensions were then prepared for imaging by putting 15 μL of cell solution onto a microscope slide with an image spacer (Grace Bio-Labs) and a #1 size coverslip (Fisher Scientific).

### Image analysis protocol

Z-stack images are first opened in Fiji, an open-source, free image analysis software [90]. The z-slice that corresponds to the midpoint of the cells is manually determined by visually inspecting the z stack. This slice is used for subsequent analysis, which is conducted in a Python (https://ipython.org) Jupyter notebook [91, 92]. For this subsequent analysis, the notebook handles opening images, segmenting cells, measuring each cell’s area and eccentricity (recorded in a data sheet), and creating PDFs These processes are described in separate subsections below, as follows:

#### Cell Segmentation

Cell segmentation is achieved using the Cellpose application programming interface (API) [36], using the ‘cyto’ model. The initial diameter for segmentation is set to 6 pixels, chosen as the smallest size observed in the image. The flow_threshold for segmentation is set to 0.4, which is the default option in the Cellpose API. The resulting masks are saved with ‘_cell_mask.tif’ as a suffix.

#### Measuring area and eccentricity

Each cell’s cross-sectional area and eccentricity are measured using the scikit-image regionprops function [93]. The size of each cell is measured as its cross-sectional cell area, found from the area covered by the mask. The shape of each cell is measured as its eccentricity, found by fitting an ellipse to each mask and calculating its eccentricity. An ellipse has two perpendicular axes with lengths a and b, a > b. Eccentricity arises from the difference in these two lengths, thus: 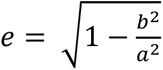. For a circle, a=b and eccentricity is zero. Values for area and eccentricity are then saved in a data sheet, where each row represents a single cell and includes the image file it originated from, along with its physical properties.

#### Generating Probability Density Functions

Kernel density estimation (KDE) is a technique used to estimate the PDF describing a set of finite data [44]. In this case, the Gaussian kernel is being used to smooth and estimate the PDF.

Mathematically, a KDE creates a PDF *f*^(*x*) using the equation,

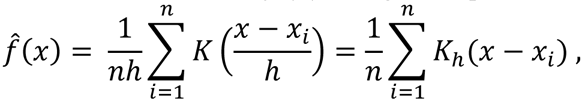

where *n* is the number of data points, ℎ is the bandwidth parameter, and *K* is the kernel function [44]. In this case, because two geometric properties are being measured, *x*_*i*_ is a two-dimensional vector containing a measured value for area and eccentricity for each cell. The Gaussian kernel function is given by,

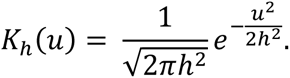

Thus, the final Gaussian kernel density estimator is,

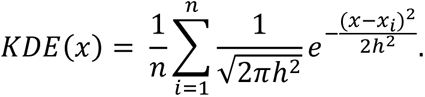

The bandwidth parameter controls how much smoothing is applied to the data: a larger bandwidth results in a smoother estimate, while a smaller bandwidth gives a more detailed but potentially noisier estimate. The PDF was calculated using the guassian_KDE function in the scipy.stats library in Python [94], with the bandwidth being chosen using the Scott’s rule of thumb,

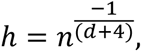

where *n* is the number of data points and *d* is the number of dimensions (here, two) [44]. The result of the KDE function is a two-dimensional normalized PDF that describes the bacterial population in terms of cell eccentricity and cell area.

We used this approach to generate PDFs for each micrograph image analyzed.

All images used and all PDFs generated are available through the Texas Data Repository.

### Measuring morphological difference in population

We created PDFs for each culture by averaging together all the PDFs obtained from images of that culture. To quantify the morphological difference between cultures, we conducted an element-wise subtraction of the PDFs for each pair of cultures. To ensure that the differences were not influenced by the direction of the subtraction, we took the absolute value of the resulting distribution. The population morphological difference was then defined as the integral of this absolute distribution. These values are shown, for every pair of cultures, in Supplementary Figure S3.

### Making the Convolutional Neural Network

The CNN was implemented using the TensorFlow [81, 82] and Keras [95] frameworks in a Keras Sequential model [95]. The model architecture consisted of several layers: the input layer was set to a 200 x 200 array, followed by a convolutional layer with 32 3×3 kernels and a Rectified Linear Unit (ReLU) activation function [39, 96]. This was followed by a max pooling layer with a 2×2 pooling kernel. Subsequently, another convolutional layer was added with 64 3×3 kernels and a ReLU activation function, followed by another max pooling layer with a 2×2 kernel. The final convolutional layer had 64 3×3 kernels with a ReLU activation function. The output of these convolutional layers was flattened and passed into a dense layer with 64 nodes and a ReLU activation function. The final output layer differed for the two models: for the Shannon Index predictor model, it consisted of 1 densely connected node with a MSE loss function, while for the top taxonomic class classifier model, it consisted of 4 densely connected nodes with a Sparse Categorical Cross Entropy loss function. Both models were optimized using the Adam optimization algorithm [69] with a learning rate of 0.001.

### Training Convolutional Neural Networks to predict taxonomic properties

The CNN we create here to predict and classify taxonomic diversity is composed of six layer types: input layer, convolutional layer, max pooling layer, flattening layer, dense layer, and output layer (Figure 7). This approach was inspired by TensorFlow’s CNN tutorial [81, 82]. The input layer is a normalized PDF “image” in the parameter space defined by cell area and cell eccentricity; this “image” is a 200 by 200 array.

The convolutional layer extracts features from the input PDF by performing a convolution operation using a trainable 3 × 3 kernel. The convolutional operation takes the dot product of the kernel and a 3 × 3 area around each element of the PDF array. The convolutional layer also has the Rectified Linear Unit (ReLU) function as an activation function. The ReLU function is defined as R(z) = max(0, z), so that it passes only positive outputs. The main purpose of using the ReLU activation function is to introduce nonlinearity into the network [39]. This nonlinearity allows the network to learn complex patterns in the data that would be difficult to capture with just linear operations. The ReLU function is used as a recommended default activation function in most modern deep learning models [81, 96–98]. The models created for this study contain 3 convolutional layers (as shown in Figure 7). The first layer contains 32 trainable kernels, the second contains 64 trainable kernels, and the third contains 64 trainable kernels.

The max pooling layer [39] reduces the dimensionality of its input by a factor of two in each dimension, by taking each 2 × 2 area in the input and returning only the maximum pixel value, effectively down-sampling the data. The max pooling layer plays a crucial role in CNNs by performing two key functions. One function is to reduce the spatial dimensions of the input data, which helps in controlling the complexity of the model and prevents overfitting [39]. The other function is to provide a degree of translation invariance to the network [39], so that the network can still recognize features in the input data even if they are shifted or translated slightly. By taking the maximum value from each local region, the max pooling layer ensures that the presence of a feature is captured regardless of its exact location within that region. The models in this study contain two max pooling layers: one after each of the first two convolutional layers.

The flattening layer, unlike the trainable layers, is not involved in parameter learning. Its role is to convert the input data into a one-dimensional array. This conversion is achieved by sequentially taking each row from the output of the preceding convolutional layer and concatenating them into a single one-dimensional array. In our models the flattening layer comes after the third convolutional layer.

The dense layer is made up of nodes that are fully connected, meaning each node receives input from every node in the previous layer [39]. Inside each node, the inputs are combined and then passed through the ReLU activation function to determine if the information should be passed along. The models only contain one dense layer of 64 nodes between the flattening and output layers. The output layer is also fully connected, with the number of nodes matching the desired output. In our research, two models were created. The model to predict the Shannon index has one output node. The classifier model to identify the top taxonomic class has four nodes, each corresponding to a primary bacterial taxonomic class.

### Model for predicting top taxonomic class

The CNN to predict the top taxonomic class represented in a sample has an output layer of 4 nodes, where each node signifies a predicted class. The output of each node is a score that the model uses to predict the top class represented by that node’s output. For example, if nodes 1, 2, 3, and 4 represent *Actinobacteria*, *Alphaproteobacteria*, *Bacilli*, and *Gammaproteobacteria*, respectively, and if the outputs of the nodes are 24, 1, 5, and 0 respectively, then the model would predict that the population had a top class representation of *Actinobacteria*.

### Model training and validation

420 PDFs were used to train and validate the CNN models. The model training process involved splitting the dataset of 420 PDFs into training and testing sets using a 70/30 ratio. This means that 70% of the PDFs were used for training the model, while the remaining 30% were reserved for testing its performance. During training, the model underwent 70 epochs, where an epoch refers to one complete pass through the entire training dataset.

To validate the model and assess its robustness, we employed repeated k-fold cross validation, or repeated random subsampling cross validation [77], with k = 10. It is a technique of retraining the model in which a new randomly chosen train-test sample split is selected with each iteration. This robust process helps to provide a more reliable estimate of the model’s performance.

After validation, the average and standard error of metrics such as MSE and accuracy were calculated and reported. MSE is a measure of the average squared difference between the predicted values and the actual values, providing an indication of the model’s predictive accuracy. The average and standard error of these metrics help to quantify the variability and reliability of the model’s performance across different iterations of the cross-validation process.

Models were trained and tested on a 2023 MacBook Pro with 16 GB of memory, and an Apple M2 Pro processor.

